# Safe and effective two-in-one replicon-and-VLP minispike vaccine for COVID-19

**DOI:** 10.1101/2020.10.02.324046

**Authors:** Alexandru A. Hennrich, Dominic H. Banda, Martina Oberhuber, Anika Schopf, Verena Pfaffinger, Kevin Wittwer, Bevan Sawatsky, Christiane Riedel, Christian K. Pfaller, Karl-Klaus Conzelmann

## Abstract

The large SARS-CoV-2 spike (S) protein is the main target of current COVID-19 vaccine candidates but can induce non-neutralizing antibodies, which may cause vaccination-induced complications or enhancement of COVID-19 disease. Besides, encoding of a functional S in replication-competent virus vector vaccines may result in the emergence of viruses with altered or expanded tropism. Here, we have developed a safe single round rhabdovirus replicon vaccine platform for enhanced presentation of the S receptor-binding domain (RBD). Structure-guided design was employed to build a chimeric minispike comprising the globular RBD linked to a transmembrane stem-anchor sequence derived from rabies virus (RABV) glycoprotein (G). Vesicular stomatitis virus (VSV) and RABV replicons encoding the minispike not only allowed expression of the antigen at the cell surface but also incorporation into the envelope of secreted non-infectious particles, thus combining classic vector-driven antigen expression and particulate virus-like particle (VLP) presentation. A single dose of a prototype replicon vaccine, VSVΔG-minispike-eGFP (G), stimulated high titers of SARS-CoV-2 neutralizing antibodies in mice, equivalent to those found in COVID-19 patients. Boost immunization with the identical replicon further enhanced neutralizing activity. These results demonstrate that rhabdovirus minispike replicons represent effective and safe alternatives to vaccination approaches using replication-competent viruses and/or the entire S antigen.

**Highlights:**

- SARS-CoV-2 S RBD antigen is preferred over entire S to preclude potential disease enhancing antibodies
- construction of a chimeric rhabdovirus minispike protein presenting RBD in native conformation
- construction of single round VSV and rabies virus replicon vaccines
- presentation of minispike antigen on cells and on noninfectious VLPs
- strong induction of SARS-CoV-2 neutralizing antibodies by the VSV replicon/VLP system in vaccinated mice

## Introduction

The current COVID-19 pandemic, caused by SARS-CoV-2, represents an exceptional challenge for our society, economy, and science. Because of the high morbidity and mortality in risk groups and possible long-term multi-organ sequelae in recovered patients regardless of age, strategies to achieve sufficient natural herd immunity are not acceptable. We are therefore witnessing unprecedented efforts and pressure to hastily develop and approve vaccines for application in a huge portion of humanity. Negligence in testing vaccine safety not only puts patients at risk but may also damage public confidence in vaccines and science in general (Thorp, 2020).

Of particular concern for vaccine safety are potentially hazardous delivery vehicles, including newly developed replicating viruses as well as harmful immune responses to inadequate antigens, known as antibody-dependent enhancement (ADE). Of special concern in case of respiratory viruses like SARS-CoV-2 is vaccine-associated enhanced respiratory disease (VAERD) (Cloutier et al., 2020; Jeyanathan et al., 2020; Lee et al., 2020), which happened previously after vaccination with conformationally incorrect viral antigens of respiratory syncytial virus (RSV). Especially, VAERD was associated with high levels of non-neutralizing antibodies. A combination of immune complex deposition, complement activation, and Th2-biased immune response led to enhancement of respiratory symptoms (Acosta et al., 2015; Polack et al., 2002; Ruckwardt et al., 2019).

The pandemic SARS-CoV-2 is a novel betacoronavirus (Wu, F. et al., 2020; Zhou et al., 2020; Zhu et al., 2020), closely related to the severe acute respiratory syndrome (SARS) virus (now named SARS-CoV-1) which emerged in 2003 (Drosten et al., 2003; Ksiazek et al., 2003). Previous work on SARS-CoV-1 was highly instructive and provided valuable blueprints for the development of the current COVID-19 vaccine candidates. In particular, Buchholz and colleagues showed that the viral surface spike (S) protein is the only virus protein that stimulates virus neutralizing antibodies (VNAs) (Buchholz et al., 2004; Bukreyev et al., 2004), which are crucial for most vaccine approaches. Accordingly, S is the main target of current COVID-19 vaccine candidates (WHO, 2020).

The class I transmembrane protein S is the primary determinant of coronavirus tropism and transmission. The S precursor protein is processed by cellular proteases into the mature N-terminal S1 and the membrane-bound S2 subunits (Bestle et al., 2020; Hoffmann et al., 2020a; Hoffmann et al., 2020b; Jaimes et al., 2020). S1 contains the receptor-binding domain (RBD) responsible for attachment of the virus to the main cellular receptor, angiotensin-converting enzyme 2 (ACE2) (Lan et al., 2020; Wrapp et al., 2020b). Binding of RBD to the receptor results in profound structural rearrangements required for membrane fusion by the S2 subunit, and release of the viral RNA genome into the cytoplasm. Molecular differences to SARS-CoV-1 S include a higher binding affinity of the RBD to the ACE2 molecule (Lan et al., 2020; Tai et al., 2020; Wrapp et al., 2020b) and the presence of a multibasic cleavage site, probably promoting proteolytic maturation and transport of the protein (Bestle et al., 2020; Hoffmann et al., 2020b; Wu, F. et al., 2020). These factors likely contribute to an extended host and organ range and the high contagiousness of SARS-CoV-2 (Kim et al., 2020; Rockx et al., 2020; Shi et al., 2020).

As accumulating data show, COVID-19 patients readily develop high antibody levels directed against the entire S protein, most of which, however, do not neutralize virus infectivity. In contrast, the overwhelming amount of RBD-binding antibodies exhibit neutralizing activity (Brouwer et al., 2020; Cao et al., 2020; Kreer et al., 2020; Premkumar et al., 2020; Zost et al., 2020). Of note, non-neutralizing antibody epitopes of SARS-CoV-1 S were previously found to enhance virus infection (Wang et al., 2016) and it was suggested that anti-S IgG from severely ill COVID-19 patients may promote hyper-inflammatory responses (Hoepel et al., 2020). At this early stage of knowledge, it therefore appears advisable to focus on the RBD immunogen in order to elicit potent neutralizing antibodies and to avoid unnecessary or potentially harmful non-neutralizing S antibodies.

Recombinant negative strand RNA viruses including rhabdoviruses like the animal pathogen VSV (Lawson et al., 1995) or the zoonotic rabies virus (Schnell et al., 1994) are attractive platforms for experimental vaccines against many emerging and neglected viral diseases, as well as for oncolytic immune therapies (for recent reviews see (Melzer et al., 2017; Zemp et al., 2018)). Rhabdoviruses are bullet shaped, cytoplasmic, and non-integrating RNA viruses encoding a single glycoprotein (G) responsible for receptor attachment and infection of cells. As illustrated before, VSV full-length or G gene-deficient (VSVΔG) vectors expressing functional S of SARS-CoV-1 induced a protective immune response in animal models (Kapadia et al., 2005; Kapadia et al., 2008). As residual pathogenicity of recombinant full length VSV is largely attributed to the glycoprotein G (Roberts et al., 1999), one strategy to attenuate VSV vaccines is the replacement of the G gene by those of heterologous envelope proteins, as exemplified in the recently approved Ebola vaccine VSV-Zebov (Ervebo®) (Matz et al., 2019). Not surprisingly, G-deficient VSV expressing fully functional SARS-CoV-2 S proteins have now been prepared and proposed as COVID-19 vaccine candidates (Case et al., 2020b; Dieterle et al., 2020; Puelles et al., 2020; Schmidt et al., 2020; Yahalom-Ronen et al., 2020; Zang et al., 2020). Importantly, and in contrast to SARS-CoV-1 S (Fukushi et al., 2006; Kapadia et al., 2005; Kapadia et al., 2008), the authentic SARS-CoV-2 spike protein can readily mediate spread and amplification of S surrogate VSVs in cell culture, organoids, and animals. Moreover, VSVΔG-SARS-CoV-2 S rapidly developed mutations in the S gene to adapt to cell culture conditions and to yield high titer viruses, as well as antibody escape mutations (Case et al., 2020b; Weisblum et al., 2020; Yahalom-Ronen et al., 2020). As attenuation of VSV evidently depends on the glycoprotein used for construction of surrogate viruses and its tropism (van den Pol et al., 2017), extensive preclinical testing is required - as was done in the case of VSV-Zebov (for review see) (Wong, 2018, Matz, 2018) - to inspire confidence in any replicating VSV surrogate virus vaccine.

Here we propose a safe and effective alternative to both replication competent viruses and expression of the full-length SARS-CoV-2 S antigen to prevent detrimental immune responses. Using structure-guided design, we developed a chimeric transmembrane RBD construct, termed “minispike”, for enhanced and structurally correct antigen presentation. In the minispike construct, the RBD domain is fused to the transmembrane stem-anchor of the G protein of rabies rhabdovirus (RABV), to allow effective expression as a cell-membrane-bound immunogen. In addition, expression of the minispike from spreading-deficient (G-deficient) VSV or RABV replicon vectors results in the secretion of non-infectious VLPs decorated with the minispike antigen. In a proof-of-principle experiment, we show that immunization with a single dose of a G-complemented VSV replicon encoding a single copy of the RBD minispike gene (VSVΔG-minispike-eGFP) is sufficient to induce high titers of SARS-CoV-2 neutralizing antibodies in mice, equivalent to those of COVID-19 patients. As the minispike is compatible with RABV, VSV and probably other rhabdoviruses, which are amenable to envelope switching, the rhabdovirus minispike system offers attractive options for a diversity of prime/boost regimens, including oral immunization with RABV G complemented viruses.

## Results

### Design of a rhabdovirus RBD-minispike

The RBD of SARS-CoV-2 spike protein was identified by sequence homology to the SARS-CoV-1 RBD and by functional studies (Lan et al., 2020; Shang et al., 2020; Tai et al., 2020; Yan et al., 2020). Structural analyses revealed a single, discrete, globular-shaped domain, able to switch between “up” and “down” configurations in the context of the pre-fusion form of the S protein, and with the up-conformation needed to engage the ACE2 receptor (Walls et al., 2020; Wrapp et al., 2020b). Based on the structure analysis we selected residues 314-541 (QTSN…KCVNF) to be included in a chimeric transmembrane minispike in which the RBD domain is presented in a natural conformation. In addition, the minispike was designed to be compatible for presentation on the cell membrane as well as for incorporation into the envelope of rhabdovirus-like particles, including VSV and RABV (Fig. 1A).

**Fig. 1.**
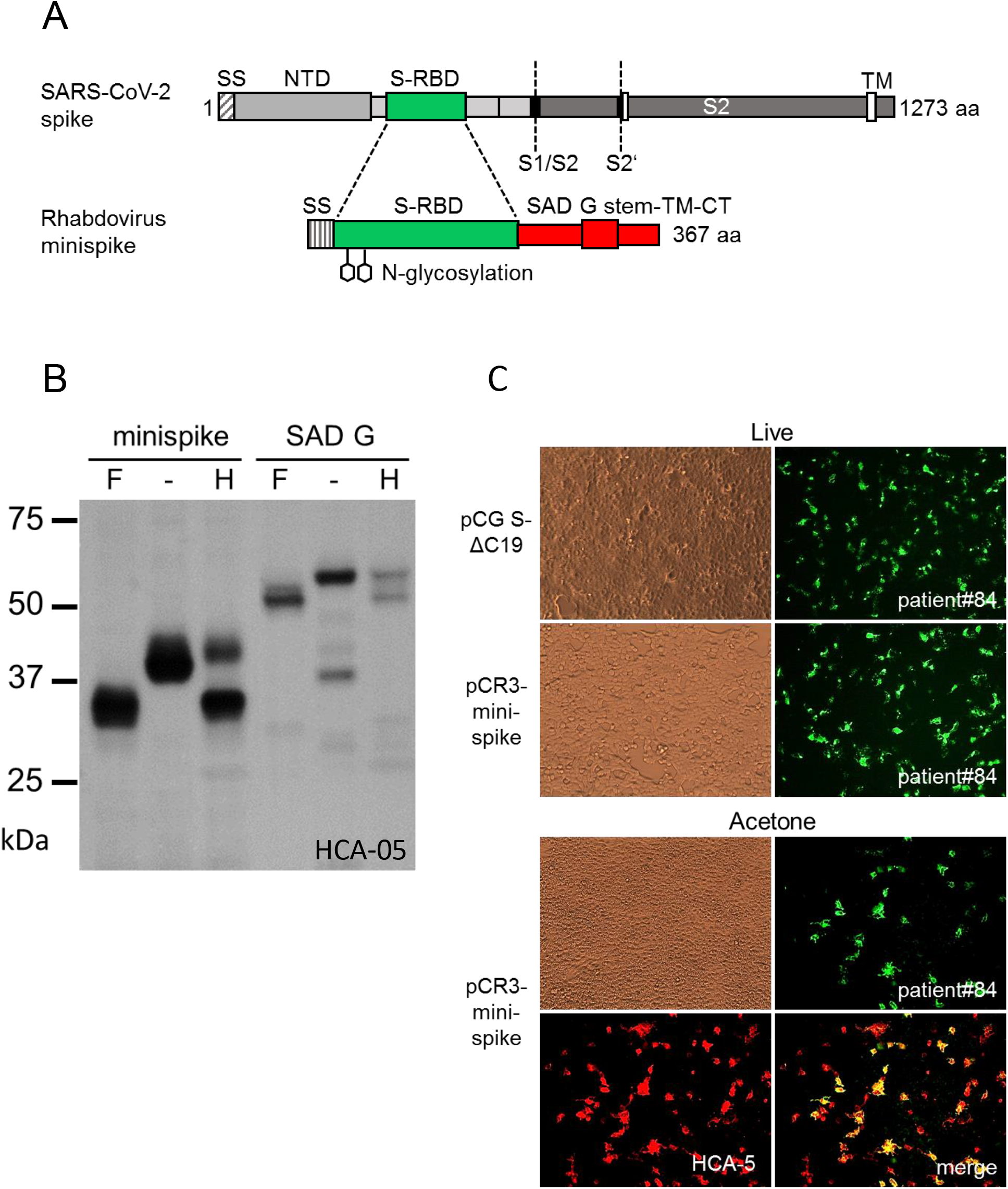
Design and expression of minispike. (A) Schematic representation of the SARS-CoV-2 spike protein (adapted from Wrapp et al., 2020), and the chimeric rhabdovirus minispike containing a hIgG signal sequence (SS) the SARS-CoV-2 RBD (green), and the RABV G stem/anchor sequence (red). Two consensus N-gylcosylation sites are indicated. S2′, S2′ protease cleavage sites, TM transmembrane domain, NTD N-terminal domain, CT cytoplasmic tail. (B) Complex N glycosylation of minispike protein. Extracts from HEK293T cells transfected with pCR3-minispike or pCAGGs-RABV-G were treated with PNGase F (+F), which cleaves off all N-linked oligosaccharides, left untreated (−) or treated with Endoglycosidase H (+H), unable to cleave complex sugars. Both minispike and RABV G acquire complex sugars, indicating transport through the Golgi apparatus. Minispike and G proteins were visualized by incubation with HCA-5 serum, recognizing the common C-tail. (C) Surface expression of minispike as revealed by staining of live HEK293T cells transfected with pCR3-minispike or pCG SΔC19, a C-terminally truncated SARS-CoV-2 S protein, with a representative patient serum 1:400 (green). Acetone-fixed and permeabilized cells expressing minispike were in addition stained with HCA-5 recognizing the RABV-derived stem-anchor (red).

The amino-terminal signal peptide from human IgG heavy chain (*Ig G HV 3-13*) was used to promote translation into the endoplasmic reticulum. The carboxy-terminus of the RBD sequence was fused via a short synthetic linker to a transmembrane stem-anchor derived from the glycoprotein of the RABV strain SAD, containing the membrane proximal part of the G ectodomain (stem), the trans-membrane domain, and the cytoplasmic sequence of SAD G (Klingen et al., 2008). The entire construct comprises 367 amino acid residues, including the signal sequence, and two N-glycosylation sites in the RBD part (<u>NIT</u>NLCPFGEVF<u>NAT</u>). The SAD G stem was selected because it should allow incorporation into the envelopes of not only RABV, but also of non-RABV rhabdoviruses, such as VSV, which has less stringent sequence requirements for membrane protein incorporation (Mebatsion et al., 1997; Schnell et al., 1998). In the case of VSV, the heterologous stem-anchor was predicted not to critically compete with VSV G incorporation needed during production of infectious single cycle VSV replicon viruses.

Expression of the minispike construct in HEK293T cells after transfection with plasmid-encoded minispike (pCR3-minispike) was at first analyzed by Western blot with an anti-SAD C-tail peptide serum (HCA-5) recognizing the RABV-derived anchor sequence (Fig. 1B). Minispike proteins were of the predicted molecular weight range, and deglycosylation experiments with PNGase F and Endo-H confirmed the presence of complex sugar chains, indicating correct processing and transport through the Golgi apparatus. Expression at the cell surface was further demonstrated by microscopic imaging and positive staining of unfixed live cells with serum from convalescent COVID-19 patients (Fig. 1C).

### Construction of minispike-expressing rhabdoviruses

Molecular clones of the Indiana strain of VSV (VSIV) (Lawson et al., 1995) were used as a basis for generation of a series of G gene-deleted VSV replicons (VSVΔG) encoding the minispike (Fig. 2A). The constructs included eGFP reporter viruses and viruses expressing single or multiple copies of the minispike gene inserted either upstream of the L gene, or at the 3’ proximal gene position, which in rhabdoviruses is transcribed most abundantly (Conzelmann, 1998; Flanagan et al., 2000). Recombinant viruses were rescued in HEK293T cells and propagated in cells transfected with VSV G plasmids or in a cell line expressing VSV G (BHK-G43) (Hanika et al., 2005). All VSVΔG viruses reached comparable titers in the range of 5×10^7^ to 3×10^8^ ffu/mL after 20-24 h of infection. G gene-deficient RABV cDNA and replicons were generated on the basis of SADΔG-eGFP and grown as described before (Buchholz et al., 1999; Ghanem and Conzelmann, 2016; Ghanem et al., 2012; Mebatsion et al., 1997).

**Fig. 2.**
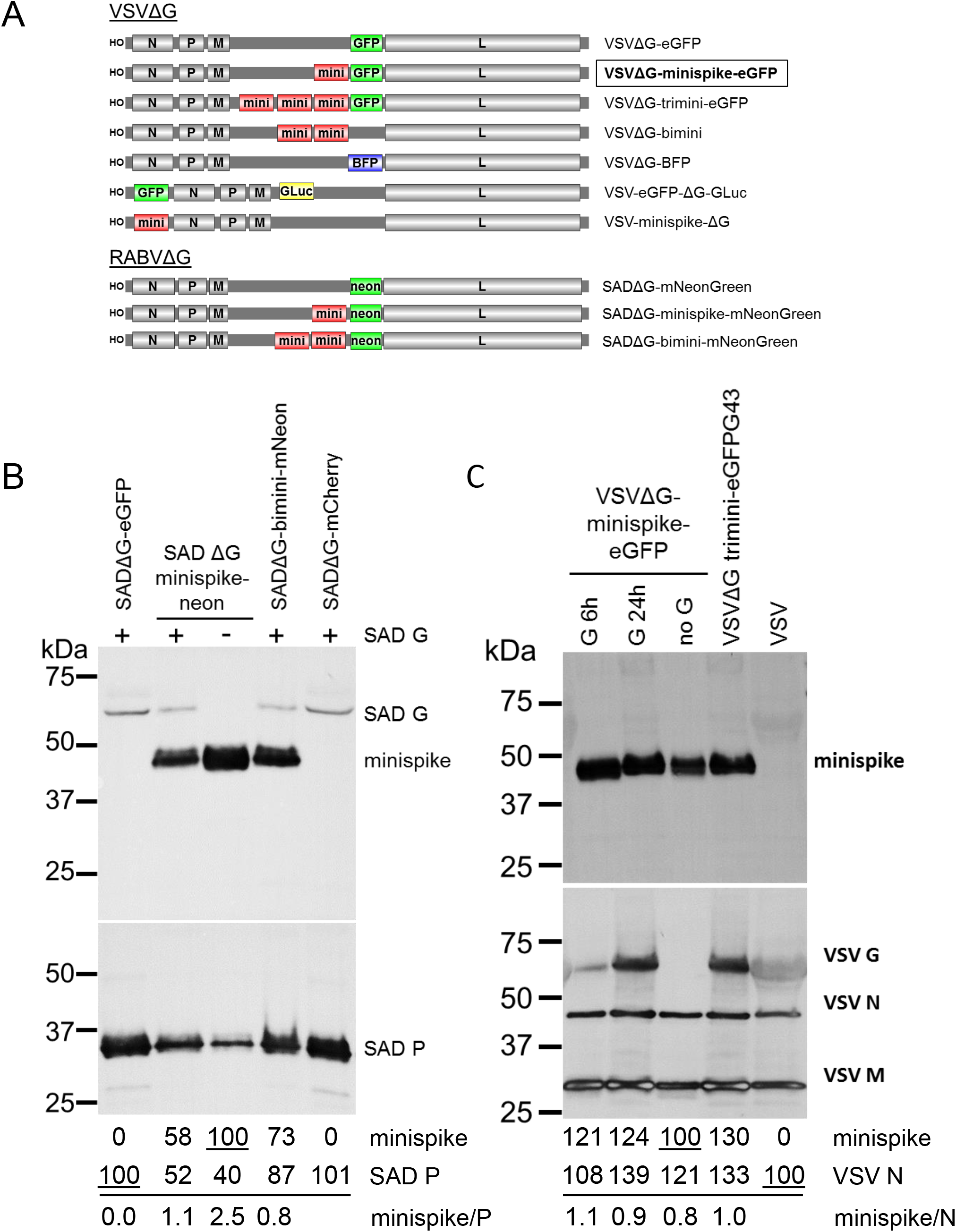
Characterization of minispike rhabdoviruses. (A) Schematic of VSVΔG and RABVΔG constructs used here. (B) Incorporation of minispike in VSV envelopes. Cell-free minispike-encoding VSVΔG viruses were generated in HEK293T cells expressing VSV G from transfected pCAGGS-VSV G for 6 or 24 hrs prior to infection, or in stable BHK-G43 cells (VSVΔG-trimini-eGFP) induced at the time of infection were purified by ultracentrifugation. Lanes were loaded with 1 million infectious units of G-containing infectious viruses, and the same volume of non-infectious viruses (no G). Blots were incubated with serum HCA-5 recognizing the RABV G-derived C-tail of the minispike (upper panel) or with anti-VSV serum recognizing viral N, M, and G proteins (lower panel). Note that G24h preparation contains G vesicles (see Fig. 3). (C) Incorporation of minispike in RABV SAD envelopes. Cell-free SADΔG-virus particles as indicated were generated in HEK293T cells in the absence or presence of RABV SAD G. Lanes were loaded with 1 million infectious units each, blots were incubated with HCA-5 recognizing the cytoplasmic tails of minispike and SAD G (upper panel) and anti-RABV P serum to determine virus load (lower panel). Quantification of band intensities indicates competition of minispike and SAD G for incorporation into virions.

**Fig. 3.**
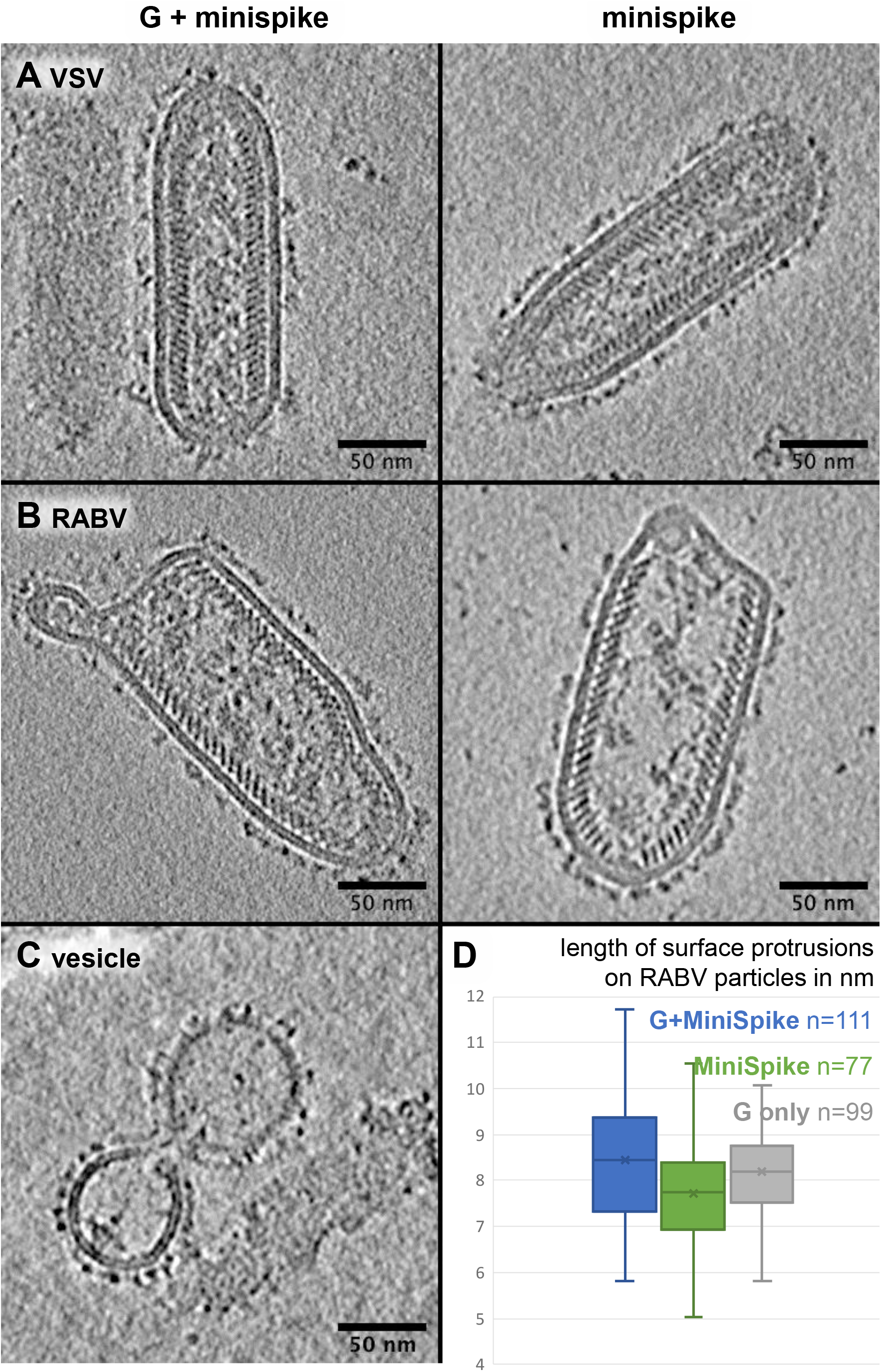
Characterization of minispike VLPs and mosaic viruses by cryo-EM. Slices through cryo electron tomograms of minispike-encoding VSV (A) and RABV (B) replicons generated in the presence of the autologous G proteins (left panel) or in the absence (right panel) indicated. Rhabdovirus particles are covered by a dense array of surface protrusions, which appears more heterogenous in the presence of G proteins. (C) VSVΔG virions produced in the presence of VSV G contain virions with mosaic envelope and G-coated non-viral vesicles (Gesicles). (D) Length distribution of surface proteins in mosaic RABV viruses (G + minispike) and particles presenting individual proteins.

### Generation of minispike VLPs and mosaic viruses

As the minispike stem-anchor is derived from the G protein of the RABV SAD strain, we first studied incorporation into virions of the autologous SADΔG-minispike-mNeonGreen and SADΔG-bimini-mNeonGreen. Supernatant virions were concentrated through a sucrose cushion by ultracentrifugation and equivalent infectious units were processed for Western blot analysis with a RABV P serum, and anti-SAD G C-tail serum to detect virus-associated minispikes and RABV G (Fig. 2B). Minispike was effectively incorporated into particles both in the absence and in presence of the parental SAD G. In the presence of SAD G less minispike was observed in RABV particles (Fig. 2B), suggesting competition of the homologous SAD G and minispike for incorporation.

To examine incorporation of the “heterologous” minispike into VSV particles, VSVΔG-minispike-eGFP stocks were produced in cells transfected with VSV G expression plasmids. For preparation of one stock, VSV G was expressed only 6 hours before VSVΔG-minispike-eGFP infection occurred, in another preparation VSV G was allowed to accumulate for 24 hours to high levels before infection. Western blot analysis of 1 million infectious units of each with anti-VSV serum revealed effective incorporation along with VSV G (Fig. 2C).

Rhabdovirus G proteins are incorporated into viral envelopes as G trimers which is driven by interaction of the C-tails with the internal M-coated viral RNP (Gaudin et al., 1992; Lyles et al., 1992; Zagouras and Rose, 1993), and incorporation supports virus budding (Mebatsion et al., 1996; Schnell et al., 1998). The presence of minispike protein in VSV envelopes could thus be due to co-incorporation with VSV G molecules as hetero-trimeric complexes. To determine whether RBD minispike alone supports budding of VSV VLPs, VSVΔG-minispike-eGFP stocks were produced in non-complementing cells and processed as above. The absence of VSV G did not prevent incorporation of minispike (Fig. 2B; lane no G), revealing autonomous incorporation and release of non-infectious minispike VSV VLPs from infected, non-complementing cells. Notably, comparable amounts of minispikes were observed in VSV particles irrespective of the presence of G (Fig. 2B). As VSVΔG viruses encoding multiple minispike genes did not show improved minispike incorporation or infectious titers, the single copy VSVΔG-minispike-eGFP was chosen for further analyses.

The composition of viral envelopes was studied in more detail by cryo-electron tomography (Fig. 3). In the absence of a rhabdovirus G protein, VSV as well as RABV VLPs contained a homogenous surface glycoprotein layer, reflecting autonomous incorporation of the minispike as suggested by the above WB experiments. The size of the globular RBD is about 60×35Å (Walls et al., 2020; Wrapp et al., 2020b), the minispike construct should thus protrude between 6 and 11 nm from the membrane. The prefusion form of rhabdovirus G protein is protruding about 8.5 nm from the virus membrane, whilst the post-fusion form is protruding about 13 nm (Albertini et al., 2012). Measuring out RABV virions expressing only G or minispike, or the combination of both, revealed differences in length of the surface protrusions (Fig. 3D). G-covered particles had surface proteins with an average length of 8.15 nm (n = 99, STD 1.07 nm) whilst in minispike VLPs this length was reduced to 7.70 nm (n = 77, STD 1.35). In the presence of both G and minispike, surface protein protrusions had an average length of 8.45 nm (n = 111, STD 1.47 nm). A direct morphological separation between G and minispike was not possible, and no higher order arrangement of the surface glycoproteins was discernible in the tomograms, suggesting random mixing.

Of note, virus preparations produced in the presence of VSV G contained non-viral vesicles with a homogenous, distinct surface protein layer. They likely represent the typical ‘Gesicles’ or G-nanovesicles formed by the autonomous budding activity of the full length VSV G protein (Mangeot et al., 2011; Rolls et al., 1994). We did not observe similar vesicular structures if RABV G or minispike were expressed. As for the parental RABV G, minispikes thus lack the ability of efficient autonomous budding.

### VSV-expressed minispike is recognized by COVID-19 patient sera

To corroborate that VSV replicons express correctly folded, processed and targeted minispike antigens that are recognized by natural antibodies of COVID-19 patients, BHK-21 cells were infected with VSVΔG-minispike-eGFP (G) and, as a control, with a VSVΔG expressing only blue fluorescent protein (VSVΔG-tagBFP) and probed with a collection of sera from patients previously tested positive for anti-S IgG in a commercial ELISA. EGFP and BFP fluorescence were used as controls to identify virus-infected cells, while bound patient IgG was detected with an Alexa555-labelled anti-human IgG secondary antibody. As illustrated in Fig. 4A for a representative serum, positive sera brightly stained VSVΔG-minispike-eGFP infected unfixed living cells, while no signal was observed with COVID-19 ELISA-negative human control sera (Fig. S1). Similarly, RABV replicon-expressed minispike was stained at the cell surface (Fig. 4B). Interestingly, while the patient sera recognized the native minispike protein as expressed by VSV and RABV replicons, they did not react effectively with reduced and SDS-denatured protein in Western blots (Fig. 4C). This indicated that the majority of the available human COVID-19 serum antibodies can only bind native conformational RBD epitopes.

**Fig. 4.**
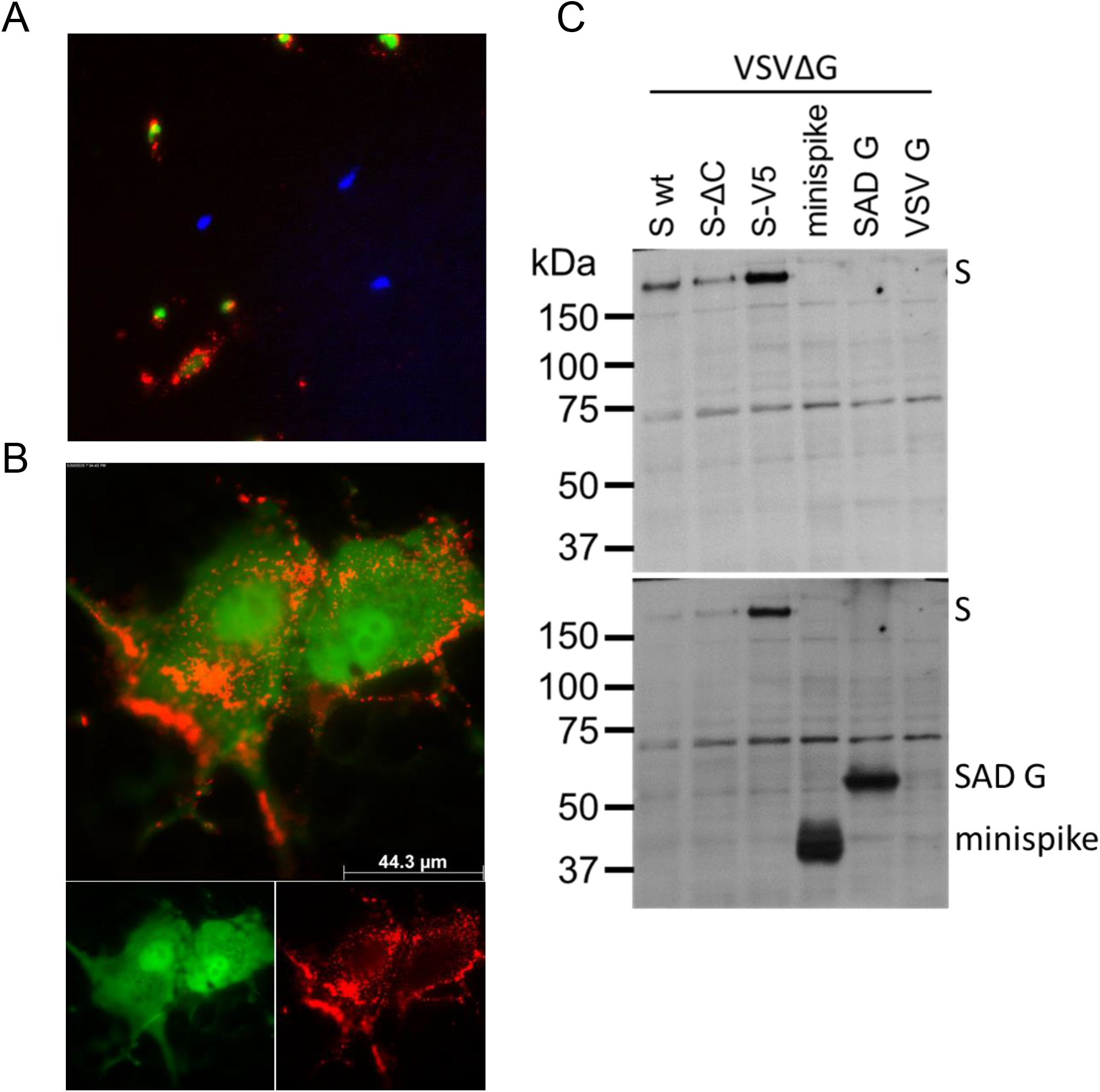
Virus minispike presents conformational RBD epitopes. (A) Cells infected with VSVΔG-minispike-eGFP (green) are recognized by IgG of S ELISA-positive patients (red), in contrast to a control virus expressing blue fluorescent protein (VSVΔG-BFP). Live unpermeabilized BHK-21 cells infected at low MOI and cultured over night at 32°C were incubated with patient serum (pat. ID 84; for other sera see Fig. S1) and stained with anti-human IgG Alexa 555, to reveal cell surface expression of minispike. 200x magnification. (B) Cells infected with a RABV replicon expressing minispike (green) are recognized by IgG of S ELISA-positive patients (red). Live unpermeabilized VeroE6 cells were infected with SADΔG-minispike-mNeongreen, incubated over night at 37°C and stained as described for (A). 1000 x magnification. (C) Poor recognition of denatured minispike protein by patient immune sera. VSVΔG-eGFP virions pseudotyped with full length wt S, a C-terminally truncated S (SΔC) or a V5-tagged S protein (S-V5) were processed for denaturing SDS Western blot and probed with a representative patient serum (Pat. #84). In contrast to the full length S proteins, denatured minispike was not readily recognized by human serum IgG. In the lower panel minispike expression was controlled by additional incubation of the same blot with HCA-5 peptide serum recognizing the C-tail present in SAD G and minispike.

In summary, the results show that the minispike protein expressed from recombinant rhabdoviruses is well recognized by conformational antibodies made in response to natural SARS-CoV-2 infection and that it thus largely mimics the conformational landscape of the natural SARS-CoV-2 S RBD. We reasoned that rhabdovirus replicons encoding chimeric minispike genes therefore represent promising and safe candidates for a COVID-19 vaccine.

### A single dose of VSVΔG-minispike-eGFP is sufficient for induction of SARS-CoV-2 neutralizing antibodies

To assess the suitability and the sufficiency of a single round VSVΔG replicon expressing a single copy of the minispike, we immunized 11-19 weeks-old BALB/c mice with VSVΔG-minispike-eGFP (G) by intra muscular (i.m.) administration. As advised by the above results, virus stocks produced under limiting (6 hrs) VSV G complementation were used, to prevent excess formation of G vesicles. Four mice received a single dose *of 1×10^6^* infectious particles, while 8 mice received an additional boost with the same virus preparation and dose 28 days following a prime vaccination. As controls, mice immunized the same way with VSVΔG-eGFP (VSV G) (n=2 for each condition) or with PBS (n=1 for each condition) were used. The 4 mice receiving only prime vaccination were sacrificed at day 28, and 4 boosted mice each at day 35 (n=4) and day 56 (n=4), to collect serum (Fig. 5A).

**Fig. 5.**
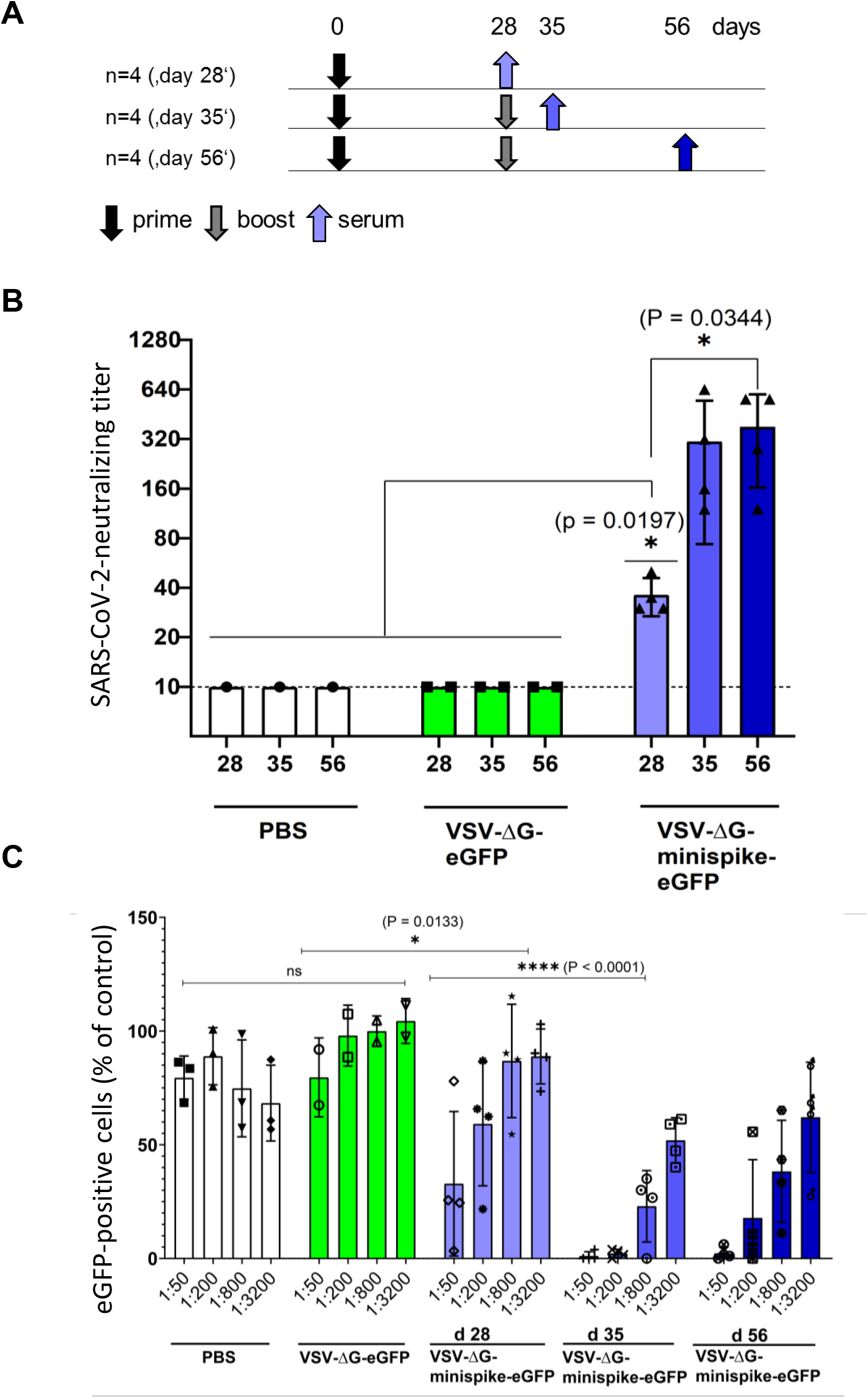
Vaccination with VSVΔG-minispike-eGFP elicits potent SARS-CoV-2 neutralizing antibodies. (A) Immunization Scheme. BALB/c mice were immunized i.m. with 1×10^6^ infectious units of VSV G-complemented VSVΔG-minispike-eGFP and controls including VSV G-complemented VSVΔG-eGFP, or PBS. Twenty-eight days after immunization serum was collected from 4 vaccinated mice, while 8 mice received an i.m. boost immunization with the same dose of virus. (B) Serum neutralization tests performed with a clinical isolate of SARS-CoV-2. The neutralizing titer of sera from vaccinated and control mice as indicated is expressed as the reciprocal of the highest dilution at which no cytopathic effect was observed. Each point represents data from one animal at the indicated time points. The bars show the mean from each group and the error bars represent standard deviations. Significant neutralizing activity was observed in mice receiving only a prime vaccination (day 28, light blue). A boost immunization further significantly enhanced neutralizing titers (days 35 and 56). (C) Neutralization of VSVΔG(S) pseudotype viruses by individual mouse sera. Mouse sera collected on day 28 (receiving prime immunization only) or at 35 and 56 days (receiving prime and boost immunization) were serially diluted as indicated and analyzed for neutralization VSV(S) pseudotype particles. GFP-encoding pseudotype virions were incubated with increasing dilutions of mouse sera or medium control before infection of VeroE6 cells. The graph shows percentage of GFP-positive cells in relation to medium controls (100%) and in dependence of dilution. Data points represent the average of three technical replicates, bars indicate standard deviation, and statistical significance was determined by one-way ANOVA.

Virus neutralization assays were performed with a SARS-CoV-2 virus isolate from Wetzlar, Germany (Bestle et al., 2020). Notably, all 4 mice immunized only once developed detectable titers of SARS-CoV-2 neutralizing antibodies in the range of 1:20-1:40 dilutions (Fig. 5B). Boost vaccination further increased neutralizing titers to 1:160-1:640.

For verification of the notable neutralizing titers in an independent assay, we also produced VSV particles pseudotyped with a functional S protein, VSV-eGFP-ΔG-GLuc (SΔC19). Neutralization assays confirmed the induction of significant levels of S-neutralizing antibodies in mice receiving a single prime vaccination and further enhancement of neutralization activity by boost immunization (Fig. 5C).

In order to directly compare the neutralizing activities of sera from vaccinated mice and from COVID-19 patients, VSV-eGFP-ΔG-GLuc (SΔC19) neutralization assays were employed. For comparison, the previously used human COVID-19 sera were used. Even sera with low ELISA IgG ratios revealed a pronounced neutralizing capacity (Fig. S2). Most intriguingly, the group of mice immunized only once (boxes labeled d 28 in Fig. 6), developed neutralizing antibodies with a capacity almost equal to those of the group of COVID-19 patients (grey boxes), illustrating a powerful induction of humoral immunity by vaccination with the single round VSVΔG-minispike-eGFP replicon. Boost immunization further enhanced neutralizing titers to exceed those of patients (Fig. 6).

**Fig. 6.**
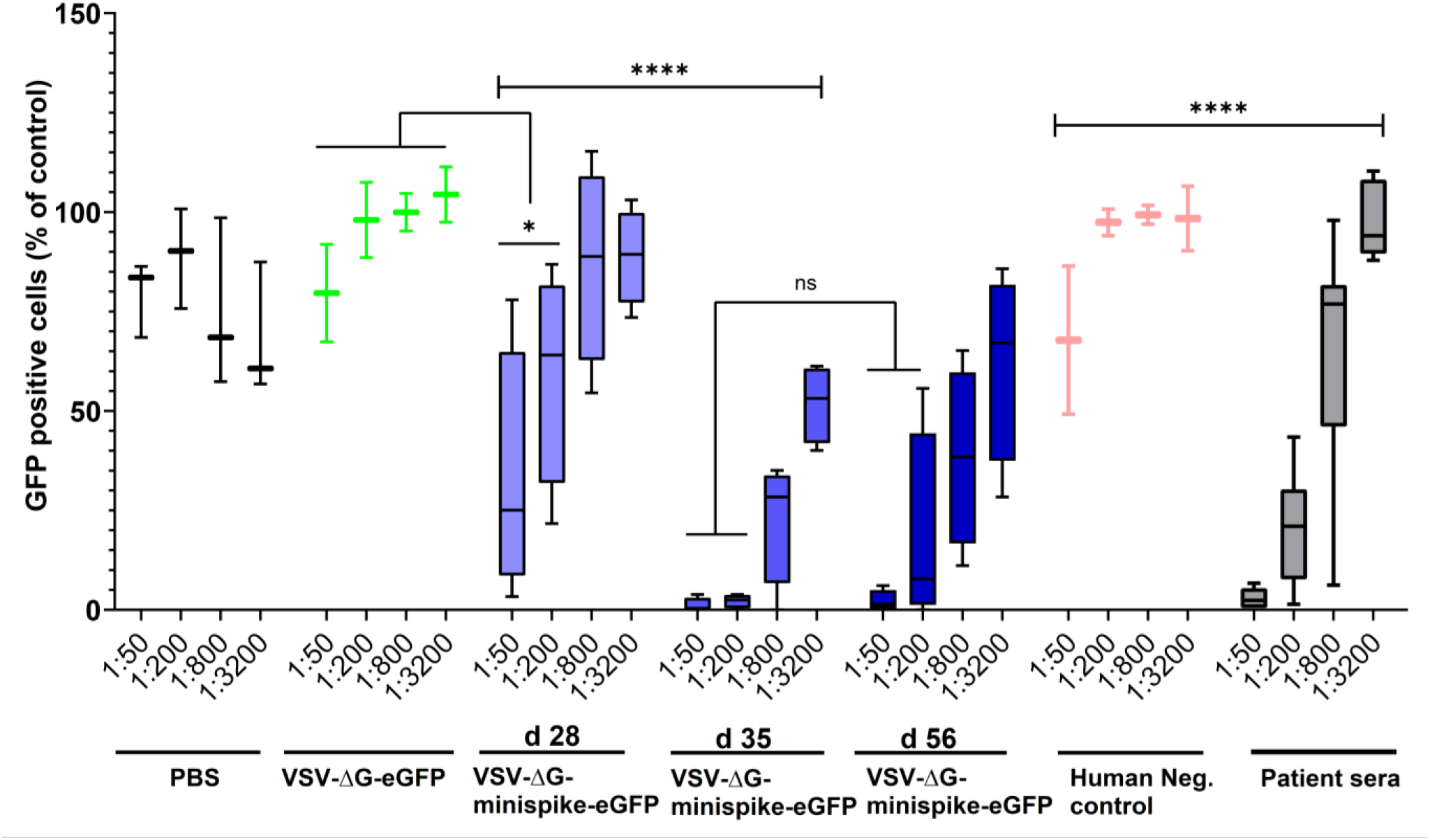
Similar virus-neutralizing titers in vaccinated mice and COVID-19 patients. VSVΔG(S) neutralization activity of sera from vaccinated mice and human immune sera tested positive for S antibodies by ELISA were compared. The graph shows percentage of GFP-positive cells in relation to medium controls and in dependence of dilution. All ELISA-positive human sera revealed VSV(S)-neutralizing activity (see Fig. S2) and are included in the grey boxes showing activity at the indicated dilutions. Primed mice (d28) exhibited neutralizing activity comparable to those of human patients, while boosted mice (d35 and d56) exhibited superior activity. Bottom and top of each box represent the first and third quartiles respectively. Whiskers represent the lowest and highest data points of the lower and upper quartile respectively. Student’s t-test and One-way ANOVA were performed to determine statistical significance.

## Discussion

There is a dangerous rush of researchers, regulatory agencies, and politicians to win the current COVID-19 vaccine race (Thorp, 2020). While it fortunately appears that SARS-CoV-2 and many of the vaccine candidates in clinical trials can readily stimulate immune responses, (see e.g. (Corbett et al., 2020; Feng et al., 2020; Mulligan et al., 2020; Suthar et al., 2020; Wu, S. et al., 2020)), stringent assessment of antigens and careful selection of delivery vehicles are still necessary before it comes to mass vaccination. Careful selection of antigens is critical for avoiding unfavorable immune reactions and disease enhancement, as in the past experienced with RSV, measles and dengue vaccines (for reviews see (Lee et al., 2020; Ruckwardt et al., 2019)). The use of viral vectors is in general associated with concerns including potential genome integration in case of all DNA vectors, high mutability of some RNA viruses, and the lack of medical control for many replication-competent live vectors. Notably, still many current candidate vaccine vectors seem to involve expression of the full-length SARS-CoV-2 S protein (WHO, 2020). In some chimeric or surrogate rhabdo- or paramyxoviruses proposed as vaccine candidates, including VSVΔG-S or Newcastle disease virus (NDV-S), functional S is even determining or extending viral spread and tropism, respectively (Case et al., 2020a; Rohaim and Munir, 2020; Sun et al., 2020a; Sun et al., 2020b; Yahalom-Ronen et al., 2020).

Here, we used a structure-guided approach to generate viral replicon vaccines meeting the requirements both in terms of virus safety and antigen harmlessness, as well as in efficacy. Our results illustrate that a small antigen, the RBD of SARS-CoV-2 if presented in the form of the present chimeric minispike protein is sufficient to induce neutralizing antibodies in mice. A single immunization with a safe, spreading-deficient single round biosafety level 1 rhabdovirus replicon was sufficient to elicit neutralizing activity similar to that of COVID-19 patients.

Assessment of antibody responses to different betacoronaviruses has recently underlined that the SARS-CoV-2 RBD is the best target for COVID-19 vaccination. While SARS-CoV-1 and MERS-CoV S proteins encode a number of VNA epitopes located outside of the RBD, the SARS-CoV-2 RBD seems to account for almost all human antibodies with potent neutralization capacity, (Barnes et al., 2020; Brouwer et al., 2020; Cao et al., 2020; Kreer et al., 2020; Zost et al., 2020), with rare exceptions (Chi et al., 2020). Furthermore, presentation of the antigen is key for the success of immunization. While this work was in progress, data on various S protein constructs became available. While soluble monomeric RBD protein suffered from limited immunogenicity, a tandem repeat single chain construct enhanced immunogenicity (Dai et al., 2020). A soluble trimeric RBD, as applied in the BNT162b1 mRNA vaccine, showed promising immunogenicity in clinical trials, including stimulation of antibodies and T cell responses (Mulligan et al., 2020; Sahin et al., 2020). In addition to trimerization, membrane anchoring seems to further improve immunogenicity, as transmembrane anchored full-length S prefusion-stabilized protein was reported to elicit higher VNA levels than corresponding secreted constructs (Corbett et al., 2020).

Here we used a previously applied strategy to present a trimeric transmembrane RBD in the form of a rhabdovirus minispike. The discrete and flexible globular structure of the RBD (Tai et al., 2020; Wrapp et al., 2020b; Yan et al., 2020) called for its combination with a rhabdovirus stem-anchor construct we have previously identified as suitable for presentation of a structurally intact protein domain (dsRED) on the surface of cells and on RABV particles (Klingen et al., 2008). The antigenic properties of the RBD in the context of the minispike remains similar to those in natural S protein, as initially indicated by binding of COVID-19 patients’ IgG to cells expressing the minispike construct (Figs 1, 4). Actually, we were initially astonished by the strong immune fluorescence signals, but in the meantime extensive characterization of natural human and animal monoclonal antibodies has revealed multiple, independent conformational epitopes in the RBD (Baum et al., 2020; Brouwer et al., 2020; Cao et al., 2020; Huo et al., 2020; Kreer et al., 2020; Tian et al., 2020; Wang et al., 2020; Wrapp et al., 2020a; Yuan et al., 2020; Zost et al., 2020). The simultaneous targeting of distinct RBD antigenic sites is of relevance not only for the efficiency of a vaccine but also in the light of potential emergence and spread of SARS-CoV-2 variants resistant against individual antibodies (Baum et al., 2020; Weisblum et al., 2020). Ongoing screening of rat monoclonal antibodies generated in response to VSVΔG-minispike-eGFP will reveal whether the chimeric minispike presents a full complement of natural RBD epitopes.

The minispike is presented copiously at the cell surface membrane, and in addition is incorporated into rhabdovirus VLPs, or mosaic viruses in the presence of G, as confirmed by immune fluorescence of cells, immune blot, and cryo-EM of virions. Reflecting the previous observations, that RABV and VSV G protein trimers are rather instable (Sissoeff et al., 2005; Zagouras and Rose, 1993), we could not immediately demonstrate a trimeric form of the minispike on the cell surface. In the context of viral envelopes, however, in which the internal RNP and matrix protein layers determine organization (Ge et al., 2010; Lyles et al., 1992; Mebatsion et al., 1996; Mebatsion et al., 1999; Riedel et al., 2019), trimeric G spikes form highly ordered paracrystalline arrays. It was suggested that the repetitive arrangement of G epitopes as observed in VSV is responsible for stimulating a very strong antibody response, by crosslinking of B cells via receptors, and possibly by contribution of T cell-independent mechanisms (Bachmann et al., 1995; Szomolanyi-Tsuda and Welsh, 1998). VLPs in general are potent immunogens, and some VLPs may be transported to local lymph nodes to promote immune responses (Roldão et al., 2010; Roy and Noad, 2009). We assume that the non-infectious minispike VLPs as generated here are synergizing with cell membrane expressed antigen, although quantification of their exact contribution to the overall immune response will require further experimentation with purified VLPs.

The excellent immunogenicity of the minispike in the context of a single-round, G-deficient VSV vaccine was illustrated by induction of high levels of SARS-CoV-2 neutralizing antibodies in mice. Notably, VNA activities equaling those of COVID-19 patients were detectable in animals receiving only a single i.m. dose of vaccine, and boost vaccination with the identical virus in the same hind leg further boosted VNA activity to levels superior to those of COVID-19 patients. VSV infection is known to induce a strong Th1 biased antiviral and anticancer immune response (Bongiorno et al., 2017; De Giovanni et al., 2020). This holds also true for VSVΔG-minispike-eGFP vaccination, as indicated by preliminary results from rats. More than 95% of S positive IgG hybridomas produced immune globulins of the IgG2 subclass, while IgG1 was only sporadically observed, thus reflecting strong Th1 immune response. A favorable humoral and cellular immune response bias as observed for BNT162b1, an adjuvanted, secreted trimeric RBD mRNA vaccine (Mulligan et al., 2020), is obtained with VSVΔG-minispike-eGFP in the absence of exogenous adjuvants.

While the chimeric minispike construct as described here appears to be immediately suitable in any genetic vaccine approach, including the auspicious mRNA platforms (Mulligan et al., 2020; Walsh et al., 2020), its full potential is accomplished in the context of the highly flexible rhabdovirus vector system, which integrates antiviral innate and adaptive immune responses. As shown here, the VSVΔG replicon complemented with little VSV G protein to mediate infection of muscle cells is highly effective in SARS-CoV-2 S RBD antigen expression after i.m. application, and intraperitoneal (i.p.) administration is supposed to be similarly effective (Kapadia et al., 2008). Boost immunizations with the same virus led to strong increase in VNA titers, revealing a moderate non-sterilizing antibody response to the VSV G protein. While results for RABVΔG-based minispike vaccines are not yet available, both VSV and RABV are amenable to envelope switching, such that pseudotyping of rhabdovirus minispike replicons with a variety of heterologous G proteins is feasible, in case repeated boosts should be necessary. RABVΔG or VSVΔG minispike vectors complemented with the G protein of widely used RABV strains like SAD in addition offer the intriguing possibility of oral immunization, in the context of prime or boost regiments.

## Methods

### Cells

HEK293T and VeroE6 (ATCC) were maintained in DMEM Medium (GIBCO) containing 10% fetal bovine serum, 1% L-Glutamine (GIBCO) and 0,5% Pen. Strep (GIBCO). BHK-G43 cells (Hanika et al., 2005), kindly provided by Georg Herrler, and BSR-MG-on cells (Finke et al., 2003) were maintained in GMEM media containing 10% fetal bovine serum, 0,5% Pen/Strep, 1% MEMs/NEAAs and 19,5mL tryptose phosphate broth (Thermo Fisher). VSV G expression in BHK-G43 cells was induced by adding 10^−9^ molar mifepristone 3 hours prior to infection, and RABV G in MG-on cells by adding 1 μM doxycycline. All cells were maintained at 37°C under 5% CO_2_.

### Construction of cDNAs

NCBI Reference Sequence NC_045512.2 of nCoV, Wuhan isolate 1, was used to synthesize human codon optimized cDNAs encoding full length HA-tagged spike (S-HA), and minispike (Thermo Fisher GeneArt). The minispike construct comprised S residues 314-541, QTSN…KCVNF fused via a GSG linker to the stem-anchor construct of SAD G described in (Klingen et al., 2008). Constructs were inserted into pCR3 for analysis of protein expression in transfected HEK293T cells and for further subcloning in RABV and VSV replicon cDNA. For production of VSVΔG(S) pseudotype viruses, we used an S-HA derived construct, pCG-S-ΔC19, kindly provided by Christian Buchholz, PEI. Plasmids for expression of wt S Protein and S variants included pCG-nCoV-S, pCG-nCoV-SΔC, and pCG-nCoV-S-V5, kindly provided by Konstantin Sparrer and Caterina Prelli Bozzo.

### Construction and rescue of recombinant rhabdoviruses

To obtain recombinant replication-competent VSVs, an infectious plasmid clone of VSIV, pVSV-eGFP (Lawson et al., 1995) (kindly provided by Jack Rose) was used to insert minispike cistrons or exchange the eGFP cassette with single or multiple copies of the minispike cistron. To yield G gene-deleted VSV replicons encoding RBD minispike the VSV G gene was replaced with minispike cassettes. To generate VSV-eGFP-ΔG-GLuc, VSVeGFPΔG (addgene #31842, kindly provided by C. Cepko) was used to insert a cistron encoding Gaussia Luciferase (GLuc) between G and L genes. pVSVdG-4BFP2 was obtained from I. Wickersham via addgene, (#64101). Virus rescue was performed in HEK293T cells transfected with the viral cDNA plasmids directing T7 RNA polymerase-driven transcription of viral antigenome (+) RNAs from a T7 promoter along with expression plasmids encoding T7 RNA polymerase and VSV helper proteins N, P, and L (pCAG-T7, -N, -P, -L; addgene #59926, #64087, #64088, #64085, respectively, all provided by I. Wickersham). Virus stocks of VSVΔG constructs were produced in HEK293T cells transfected with pCAGGs-VSV G, or BHK-G43 and concentrated by ultracentrifugation over a 20% sucrose cushion in a SW32 rotor at 24,000 rpm and 4°C for 2h.

Recombinant cDNAs of RABVΔG expressing minispike and mNeonGreen were generated by replacement of the eGFP cassette of pHH_SADΔG-eGFP_SC with two transcription units and rescued into virus in cells providing RABV N, P, L, and T7RNA polymerase as described before (Buchholz et al., 1999; Ghanem and Conzelmann, 2016; Ghanem et al., 2012). RABVΔG replicons were propagated in MG-on cells providing SAD G (Finke et al., 2003).

### Western blots

Laemmli SDS-PAGE in 6% stacking and 10% separating Bis-Tris gels and Western blot analysis on semi-dry-blotted PVDF membranes was done as previously described (Schuhmann et al., 2011). Briefly, membranes were incubated with primary antibodies at 4°C overnight, and after three times washing with TBS-T incubated for 2 hrs with horseradish peroxidase-conjugated secondary antibodies at room temperature. Bio-RAD Clarity Western Enhanced Chemiluminescence (ECL) Substrate kit was used for detection in a Fusion Fx7 imaging system.

### Microscopy

For live cell imaging, VeroE6 or HEK293T cells were seeded into poly-D-lysine (Millipore-Sigma)-coated multiwell plates one day prior to infection with VSV replicons at the indicated MOIs or plasmid transfection by lipofection, respectively. Infected cells were incubated overnight at 32°C. Minispike was detected by incubation with human COVID-19 patient sera or the serum of a healthy donor diluted 1:300 in DMEM fluorobrite for one hour at 37°C and subsequent staining with anti-Human IgG (H+L) AlexaFluor555 (1:2,000 in DMEM Fluorobrite, 1h, 37°C) and imaged after washing with DMEM fluorobrite. For fixation and permeabilization, infected or transfected cells were washed once with PBS and treated with 80% acetone for 20 min at room temperature. After blocking with 5% bovine serum albumin (BSA) in PBS for 1h at room temperature, and three times washing, cells were incubated with COVID-19 patient sera and HCA-5 rabbit peptide serum in PBS with 1% BSA over night at 4°C. The cells were then washed three times with PBS and incubated with anti-Human IgG (H+L) AlexaFluor488 and anti-Rabbit IgG (H+L) AlexaFluor555 for one hour at room temperature. After 3 washing steps, cells were imaged on a Leica DMi8 microscope equipped with LED405 (blue), GFP (green), TXR (red) and Cy5 (far red) filter cubes.

### Cryo-electron microscopy

Concentrated preparations of rhabdovirus pseudotype particles were added to glow discharged Quantifoil 200 mesh 2/1 holy carbon copper grids in the presence of Aurion protein A 10nm gold beads. Vitrification was performed either with a manual plunging unit or a FEI Vitrobot. Grids were analyzed in a FEI Glacios or a FEI Talos Arctica operated at 200kV and bidirectional or dose symmetric tilt series were acquired with a FEI Falcon 2 direct electron detector. Tomograms were subsequently reconstructed with etomo and visualizsed with 3dmod (Kremer et al., 1996).

### Animal experiments

Mouse immunization studies were carried out in the animal housing facility of the Paul-Ehrlich-Institute, in compliance with the regulations of German animal protection laws and authorized by the responsible state authority (V54-19c20/15-F107/1058). 11-19 weeks old female BALB/c mice received one or two intramuscular injections of 1×10^6^ ffu of either VSVΔG-minispike-eGFP (VSV G), or VSVΔG (VSV G) “empty” dissolved in 30 μl PBS, or an equal volume of PBS alone, four weeks apart (prime or prime/boost). Mice receiving only a single dose of vaccine were sacrificed on day 28, mice from the prime/boost group were sacrificed on days 35 or 56. The mice were anesthetized by intraperitoneal injection of 100 mg/kg body weight ketamine and 10 mg/kg body weight xylazine and exsanguinated retroorbitally or by cardiac puncture. Whole blood was collected in Z-gel containing tubes (Sarstedt). Serum was separated by centrifugation at 14,000 g for 15 min at room temperature and stored at −20°C.

### Virus neutralization assays

SARS-CoV-2 neutralizing antibody titers were determined by mixing serial dilutions of serum collected from mice at the indicated time points with 10^2^ TCID50 of SARS-CoV-2 (Wetzlar isolate), kindly provided by Eva Friebertshäuser (Bestle et al., 2020). Virus and serum dilutions were incubated at 37°C for 20 min before 50 μl of VeroE6 cells were added to each well. After incubation for 3 days at 37°C, cell monolayers were stained with PBS containing 4% paraformaldehyde (PFA) and 1% crystal violet. The neutralizing titer is expressed as the reciprocal of the highest dilution at which no cytopathic effect (CPE) was observed.

Neutralization of VSV (S) pseudotyped viruses were performed as follows: HEK293T cells transfected with pGC-SΔC19 (obtained from Christian Buchholz) for one day were infected with VSV G-complemented VSV-eGFP-ΔG-GLuc at a MOI of 1. After 3 hrs incubation, excess input virus was removed by thorough washing. After incubation for 24h, supernatant was collected, and S pseudotype virions concentrated by ultracentrifugation through a sucrose cushion, resuspended in PBS and titrated on VeroE6 cells in the presence of VSV-neutralizing hybridoma supernatant (I1-Hybridoma, ATCC CRL-2700) to block residual input G-containing virus. Briefly, VeroE6 cells were seeded at a density of 1×10^4^ cells per well in 96 well-plates, and incubated with 10^2^ ffu of S pseudotype viruses in the presence of I1 supernatant, and with cell culture medium as a control, or increasing dilutions of mouse or human sera, as indicated, in a total volume of 100 μL. Infectious units were determined after one-day incubation by manual counting of fluorescent cells in triplicate experiments.

### Data representation and statistical analysis

Statistical analyses were performed using GraphPad Prism version 8.4.3. Unless otherwise stated, data are from at least three technical replicates. Statistical significance was calculated using 2-tailed Student t-test or one-way ANOVA; results are indicated in figures (* p=0.05; ** p<0.01; *** p<0.001; **** p<0.0001; ^ns^ not significant).

## Lead Contact and Materials Availability

Requests for material can be directed to the lead contact, Karl-Klaus Conzelmann. All materials and reagents will be made available upon installment of a material transfer agreement (MTA).

## Acknowledgements

We thank Konstantin Sparrer, Ulm University, for interactions of all sorts, and Yassine Haddad for help in establishing the VSV rescue system. Plasmids were kindly provided by John K. Rose (VSIV cDNAs), Ian Wickersham and Conny Cepko (addgene VSV cDNAs), Christian Buchholz (SΔC19), Konstantin Sparrer, Caterina Prelli Bozzo, and Stefan Pöhlmann (S, S-ΔC, S-V5). We thank Georg Herrler for providing BHK-G43 cells, Ralf Bartenschlager for ACE2-expressing cells, and Eva Friebertshäuser and Stephan Becker for the SARS-CoV-2 isolate. Max Münchhoff and Patricia Spaeth kindly provided patient sera. Research in the authors’ labs is supported by the Deutsche Forschungsgemeinschaft (DFG, German Research Foundation) - Project-ID 369799452 - TRR237 A12”, - Project-ID 118803580 - SFB 870 Z1”, and Co260/5, and by FöFoLe (Förderprogramm für Forschung und Lehre) of the Medical Faculty of the LMU Munich (FöFoLe M 09/2017). CKP is supported by the DFG Collaborative Research Center 1021 (project number 197785619/B09) and by intramural funding of the German Federal Ministry of Health (BMG), and BS by the German Center for Infection Research (DZIF), project number HZI2016Z9. Cryo-electron microscopy was performed at the Electron Microscopy Facility at Vienna BioCenter (VBC) Core Facilities (VBCF), Austria, and at the CEITEC. The help of Thomas Heuser and Jiří Nováček with the cryo-EM data collection is greatly appreciated. CR was supported by CIISB research infrastructure project LM2018127 and project number CIISB #200084C funded by MEYS. We thank Konstantin Sparrer and Norbert Tautz for expert review of the manuscript.

## Author contributions

AAH, KKC, and CKP contributed to the concept of the study, AAH designed and cloned viruses. CR performed cryo-EM analyses, KW, BS and CKP performed mouse experiments and BS performed SARS-CoV-2 virus neutralization assays. DHB, MO, AS, and VP produced S pseudotype viruses and performed neutralization assays, DHB conducted statistical analysis. All authors contributed to writing and editing of the manuscript and have agreed to the submitted version.

## Supplementary Figures

**Fig. S1 (related to Fig. 4).**
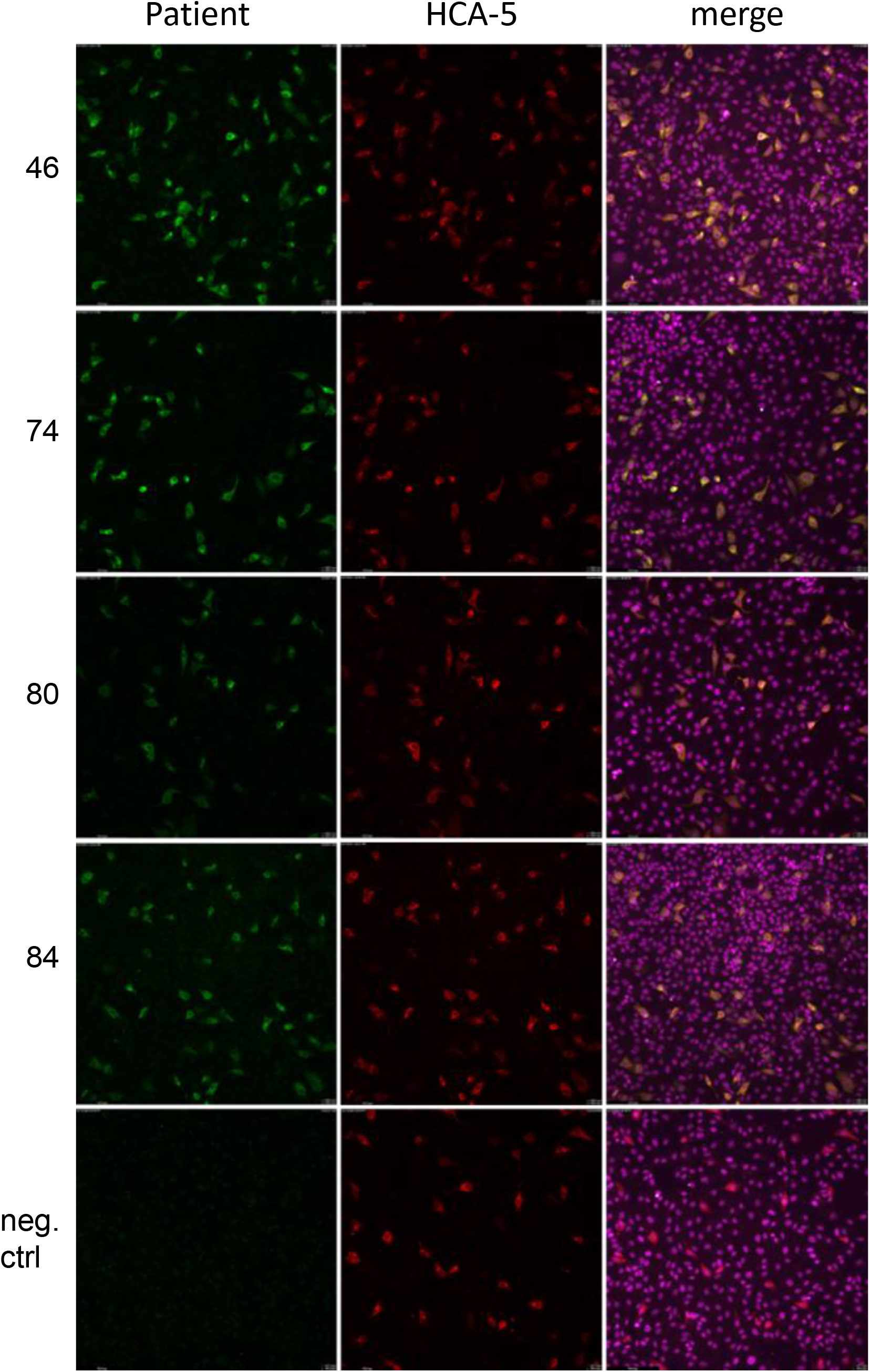
Recognition of VSV-expressed minispike by patient sera. Vero E6 cells were infected with VSV-ΔG-bimini overnight at 32°C, fixed in 4% PFA and permeabilized with 0.05% Saponine followed by incubation with different S ELISA-positive sera of COVID-19 patients, and HCA-5 as a control for minispike expression. In contrast to a human negative control serum, all patient sera stained cells expressing minispike with anti-human IgG (H+L) Alexa 488 (green). ToPro3 (magenta) was used to counterstain nuclei.

**Fig. S2. (related to Fig. 6).**
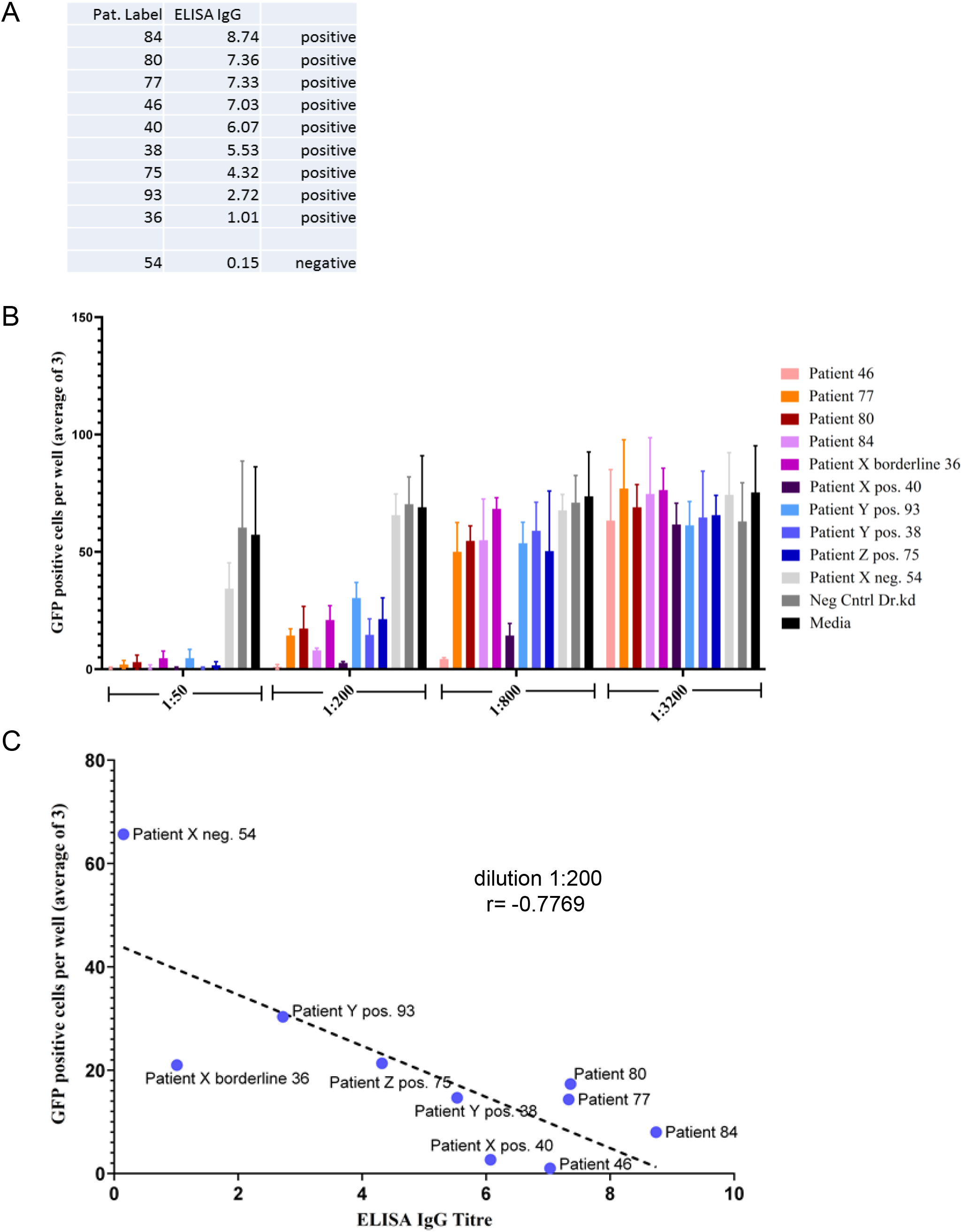
Characterization of COVID-19 patient sera. (A) ELISA IgG ratio of sera (B) VSV(S) neutralizing activity of human sera. Graph shows reduction of ffu of VSVeGFP-ΔG-GLuc S pseudotype viruses after incubation with sera at the indicated dilutions. (C) Comparison of ELISA titers and neutralizing activity at 1:200 dilution.

